# Age dependence of systemic bone loss and recovery following femur fracture in mice

**DOI:** 10.1101/291906

**Authors:** Armaun J. Emami, Chrisoula A. Toupadakis, Stephanie M. Telek, David P. Fyhrie, Clare E. Yellowley, Blaine A. Christiansen

## Abstract

The most reliable predictor of future fracture risk is a previous fracture of any kind. The etiology of this increased fracture risk is not fully known, but it is possible that fracture initiates systemic bone loss leading to greater fracture risk at all skeletal sites. In this study we investigated systemic bone loss and recovery following femoral fracture in young (3 month old) and middle-aged (12 month old) mice. Transverse femur fractures were created using a controlled impact, and whole-body bone mineral density (BMD), trabecular and cortical microstructure, bone mechanical properties, bone formation and resorption rates, mouse voluntary movement, and systemic inflammation were quantified at multiple time points post-fracture. We found that fracture led to decreased whole-body BMD in both young and middle-aged mice 2 weeks post-fracture; this bone loss was recovered by 6 weeks in young, but not middle-aged mice. Similarly, trabecular bone volume fraction (BV/TV) of the L5 vertebral body was significantly reduced in fractured mice relative to control mice 2 weeks post-fracture (−11% for young mice, −18% for middle-aged mice); this bone loss was fully recovered by 6 weeks post-fracture in young mice. At 3 days post-fracture we observed significant increases in serum levels of interleukin-6 and significant decreases in voluntary movement in fractured mice compared to control mice, with considerably greater changes in middle-aged mice than in young mice. At this time point we also observed increased osteoclast number on L5 vertebral body trabecular bone of fractured mice compared to control mice. These data show that systemic bone loss occurs after fracture in both young and middle-aged mice, and recovery from this bone loss may vary with age. This systemic response could contribute to increased future fracture risk following fracture, and these data may inform clinical treatment of fractures with respect to improving long-term skeletal health.

## Introduction

Many risk factors for osteoporotic fractures have been identified, but one of the most reliable predictors of a future fracture is a previous fracture of any kind ^(1–7)^. Subjects who have sustained a previous fracture are several times more likely to sustain a future fracture than those with no history of a fracture, even when controlling for bone mineral density (BMD) ^(2-6,8)^. The risk of future fractures increases with the number of prior fractures ^(2,8)^, and is maintained even when the fracture occurs at an unrelated skeletal site ^(2,4,7,9)^. Increased fracture risk is not constant following an index fracture; it is highest in the first 1-2 years following index fracture, then decreases over subsequent years ^(1,7,10^, but remains higher than that of the general population. Age at first fracture is also predictive of the increased risk of future fractures. Wu et al. showed that any fracture in women (not related to motor vehicle accidents) that occurred in adulthood (age 20-50) is associated with increased risk of fracture after the age of 50, while fractures that occur before the age of 20 are not correlated with future fracture risk ^(3)^. A similar study by Amin et al. showed that distal forearm fractures in childhood (≤18) are not predictive of future fracture risk in women, but are highly associated with future fracture risk in men ^(9)^.

Traumatic injuries initiate a systemic inflammatory response that can lead to tissue adaptation at sites not directly affected by the initial injury ^(11,12)^. In bone, it is well established that the bone formation and bone resorption rates near a fracture are higher than baseline activity; this is known as regional acceleratory phenomenon (RAP) ^(13–15)^. Additionally, Ziegler et al. demonstrated a systemic acceleratory phenomenon (SAP) following creation of a tibial defect in rats ^(16,17)^. In this study, creation of a burr hole defect in rat tibiae resulted in increases in mineralizing surface, mineral apposition rate, and bone formation rate in the lumbar spine. These are critical observations, as a “transient osteoporosis” has been noted during RAP because of the temporal lag between resorption and formation during bone healing. It is unclear if a similar transient osteoporosis occurs systemically with fracture, and whether animals and humans are able to fully recover from this bone loss, or if this phenomenon results in weaker bones and increased risk of future fractures at all skeletal sites.

The goal of our study was to investigate changes in bone mass systemically, and bone microstructure and mechanical properties at distant skeletal sites following femur fracture in mice, to investigate potential mechanisms that may contribute to this systemic adaptation, and to determine the effect of age on this response. We hypothesized that femoral fracture in mice would initiate systemic inflammation, increased rates of bone formation and resorption, and decreased voluntary movement, ultimately resulting in long-term deficits in bone volume and strength in middle-aged (12 month old) but not young (3 month old) mice.

## Methods

### Animals

A total of 334 mice (female C57BL/6J; Envigo, Somerset, NJ) were used for this study, including 161 young (3 month old), and 173 middle-aged (12 month old) mice (Fig. 1). Fractured mice were subjected to transverse femur fracture stabilized by an intramedullary pin (described in more detail below); Sham mice were subjected to surgical placement of the intramedullary pin without fracture; Control mice were subjected to anesthesia and analgesia only. Mice were euthanized at 2, 4, or 6 weeks post-injury for micro-computed tomography and mechanical testing (n = 6-10 per treatment/age). Separate groups of mice (n = 6-10 per treatment/age) were euthanized at 3 days, 2, 4, or 6 weeks post-injury for assessment of systemic inflammation, bone formation, and bone resorption. Additional mice were sent to the UC Davis Mouse Biology Program (https://mbp.mousebiology.org/) following fracture for whole-body duel-energy x-ray absorptiometry (DXA) scanning and open field assessment voluntary movement at 4 days, 2, 4, and 6 weeks post-injury (n = 4-6 per treatment/age). Two young mice and one middle-aged mouse were euthanized due to intramedullary pin misplacement or complications after surgery. All mice were euthanized via CO_2_ asphyxiation followed by cervical dislocation. Mice were cared for in accordance with the guidelines set by the National Institutes of Health (NIH) on the care and use of laboratory animals. Mice were housed in Tecniplast conventional cages (Tecniplast SPA, Buguggiate, Italy), with Bed-o’ Cob bedding (The Andersons Inc., Maumee, OH), with 3-4 mice per cage, 12-hour light/dark cycle, 20-26° C ambient temperature. Mice had ad libitum access to food (Harlan irradiated 2918 chow) and autoclaved water, and were monitored by husbandry staff at least once a day, 7 days a week, with monthly health care checks by a veterinarian. All procedures were approved by the Institutional Animal Care and Use Committee of the University of California, Davis.

**Figure 1.**
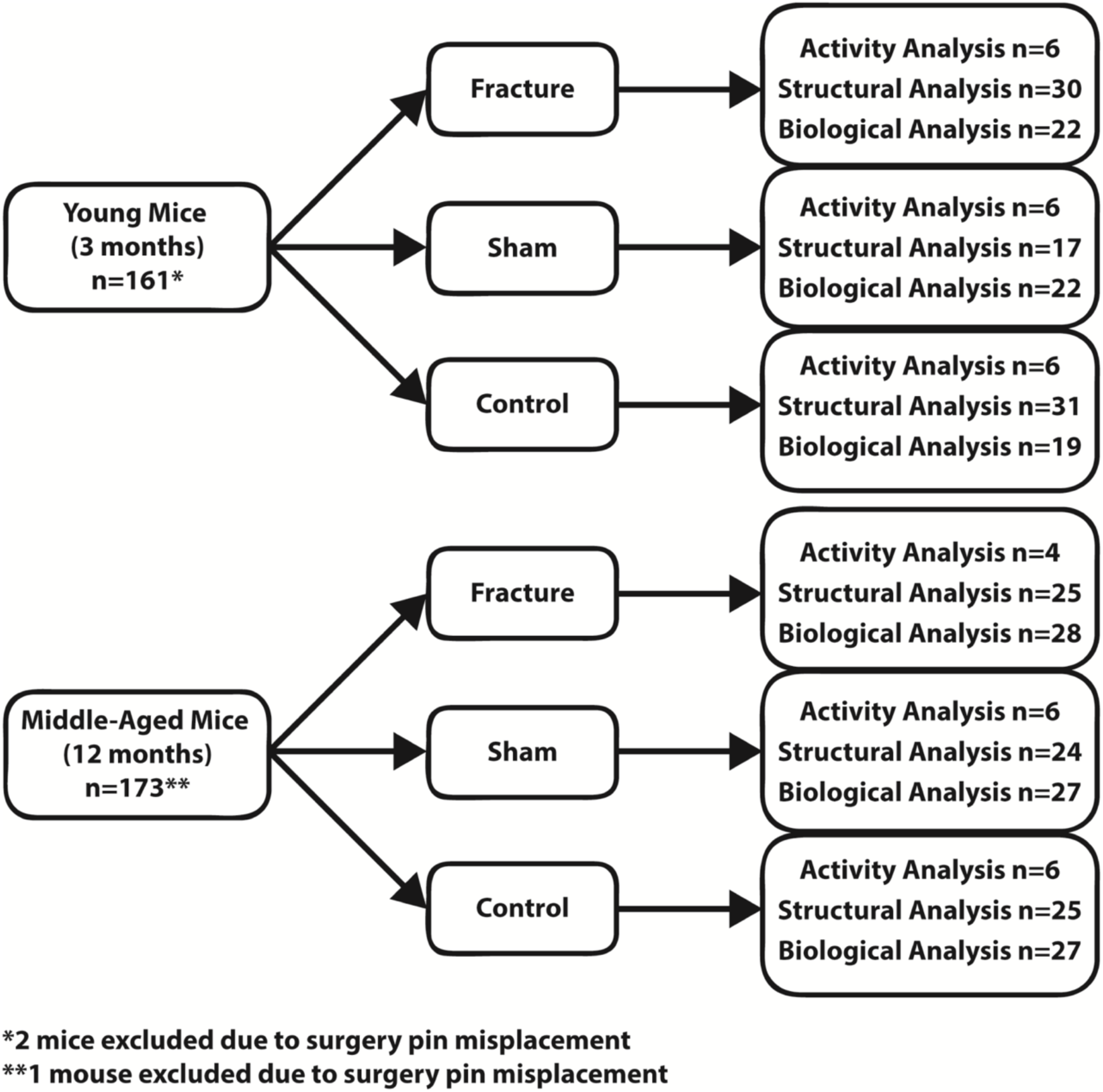
Study design of young (3 month old) and middle-aged (12 month old) mice after femoral fracture. Activity Analysis consisted of Open Field testing and DXA imaging of whole-body BMD and BMC. Structural Analysis included µCT imaging and mechanical testing of the L5 vertebral body, contralateral femur, and contralateral radius, and finite element analysis of the L5 vertebral body. Biological Analysis included dynamic histomorphometry of the contralateral tibia, TRAP staining of the L5 vertebral body, and measurement of inflammatory cytokines in serum using ELISA.

### Creation of Femur Fracture

All mice were given a pre-operative dose of buprenorphine (0.05 mg/kg) and then anesthetized with isoflurane inhalation. For Fractured and Sham mice, an incision was created on the lateral side of the right knee and a 0.01 in. stainless steel wire pin was inserted into the medullary canal as described by Bonnarens and Einhorn ^(18)^. For Fractured mice, transverse fractures were then created with a controlled lateral impact using a modified fracture apparatus ^(19,20)^. Immediately post-fracture, mice were imaged with planar radiographs (HE100/30+ X-ray machine, MinXRay, Northbrook IL) with a CXD1-31 plate (Canon, Lake Success, NY) to confirm pin positioning and a transverse mid-diaphyseal fracture. Full weight-bearing and unrestricted activity was permitted postoperatively.

### Micro-Computed Tomography Analysis of Cortical and Trabecular Bone

Following euthanasia, contralateral femora, contralateral radii, and L5 vertebrae were removed and preserved in 70% ethanol. Bones were scanned using micro-computed tomography (SCANCO Medical, µCT 35, Brüttisellen, Switzerland) with 6 µm nominal voxel size (x-ray tube potential = 55 kVp, current = 114 μA, integration time = 900 ms, number of projections = 1000/180**°**) according to the JBMR guidelines for μCT analysis of rodent bone structure ^(21)^. All analyses were performed using the manufacturer’s analysis software. Trabecular bone was analyzed at the L5 vertebral body and distal femoral metaphysis. Trabecular regions of interests were manually drawn on transverse images excluding the cortical surface. The region of interest for the L5 vertebral bodies extended from the cranial growth plate to the caudal growth plate, excluding the transverse processes. The region of interest for the distal femoral metaphysis started adjacent to the metaphyseal growth plate and extended 100 slices (600 µm) proximal. Trabecular bone volume fraction (BV/TV), trabecular thickness (Tb.Th), trabecular separation (Tb.Sp), trabecular number (Tb.N), connectivity density (Conn.Dens), and tissue mineral density (TMD) were determined. Cortical bone was analyzed at the mid-diaphysis of the femur and radius. Regions of interest included 100 slices (600 µm) centered at the longitudinal midpoint of each bone. Bone area (B.Ar), total cross-sectional area (Tt.Ar), medullary area (Me.Ar), cortical thickness (Ct.Th), maximum and minimum bending moments of inertia (I_max_ and I_min_), bone mineral content (BMC), and tissue mineral density (TMD) were determined. The cortical shell was also analyzed at the L5 vertebral body, using a region of interest that extended from the cranial growth plate to caudal growth plate excluding the trabecular compartment. Bone area (B.Ar), cortical thickness (Ct.Th), and tissue mineral density (TMD) were determined.

### Three-Point Bending Mechanical Testing

Contralateral (uninjured) femora and contralateral radii were mechanically tested in three-point bending to determine structural and material properties of cortical bone. Bones were removed from 70% ethanol and rehydrated for 10-15 minutes in phosphate buffered saline (PBS) prior to mechanical testing using a materials testing system (ELF 3200, TA Instruments, New Castle, DE) with the anterior aspect of each bone in tension. Data was collected at a sampling frequency of 50 Hz, the lower supports had a span of 7.45 mm, and the center loading platen was driven at 0.2 mm/sec until failure. Resulting force and displacement data were analyzed to determine stiffness, yield force, ultimate force, energy to fracture, and post-yield displacement. Bending moments of inertia from μCT scans were used to calculate modulus of elasticity, yield stress, and ultimate stress for each bone using Euler-Bernoulli beam theory.

### Compression Testing of L5 Vertebral Bodies

L5 vertebral bodies were mechanically tested in compression to determine structural properties of trabecular bone as previously described ^(22–24)^. Posterior elements were removed, and the endplates of the vertebrae were minimally shaved flat, resulting in a consistent sample height of 2 mm. Vertebrae were removed from 70% ethanol and rehydrated for 10-15 minutes in phosphate buffered saline (PBS) prior to mechanical testing using a materials testing system (ELF 3200, TA Instruments, New Castle, DE). Data was collected at a sampling frequency of 50 Hz, and the loading platen was driven at 0.05 mm/sec until failure. Resulting force and displacement data were analyzed to determine compressive stiffness, yield force, and ultimate force.

### Finite Element Analysis of Compression Tests for L5 Vertebrae

Finite element analysis was used to further estimate mechanical properties of L5 vertebral bodies. µCT scans of L5 vertebral bodies were uploaded into ImageJ ^(31)^. Each scan was then adjusted so the longitudinal axis of the vertebral body was perpendicular to the simulated compression plates and processed to isolate a 2 mm section of the vertebral body, including both cortical and trabecular bone. FE models were created of each vertebral body using a custom software, and analysis of FE models followed our published methods ^(25–27)^. Each model was subjected to simulated compression testing using custom linear finite element modeling software. The hard tissue in the models was assumed to be isotropic and uniform with a Poisson’s ratio of

0.5. Young’s modulus was calculated for each specimen based on mean tissue mineral density as described by Easley et al. ^(28)^ using the equation:

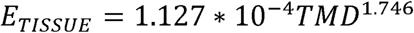

in which E_TISSUE_ is the modulus of the bone tissue (GPa) and TMD is the mean tissue mineral density (mg HA/cm^3^) determined from µCT. FE models were tested in compression; the top of the model was compressed uniformly to an apparent strain of 0.5%, and stiffness along the longitudinal axis of the vertebral body was calculated for each specimen. The top and bottom of the cylinder were constrained using fixed boundary conditions to simulate mechanical testing between glue-bonded platens.

### Quantification of Mouse Voluntary Activity Using Open Field Analysis

Following surgery, 18 young mice and 16 middle-aged mice (n = 4-6/injury/age group), were transferred to the UC Davis Mouse Biology Program for analysis of voluntary movement and activity. At each time point (day 4, weeks 2, 4, and 6 post-fracture) mice were individually housed for 24 hours in an isolated, enclosed chamber (18 inch square) designed to evaluate voluntary activity using infrared sensors (ACTIVITY SENSOR, Ohara Co. Ltd., Tokyo, Japan) as previously described ^(29)^. Measured parameters included total distance traveled, vertical sensor breaks (indicating rearing), resting time, ambulatory time, time spent in the perimeter, and time spent in the center.

### Dual Energy X-Ray Absorptiometry (DXA) Imaging of Whole-Body Bone Mass and Density

Mice transferred to the UC Davis Mouse Biology Program were imaged with DXA (Lunar PIXImus II Densitometer, GE Medical Systems, Chalfont St. Giles, UK) to quantify longitudinal changes in whole-body bone mass and density. At each time point (day 4, weeks 2, 4, and 6 post-fracture), mice were anesthetized using isoflurane inhalation and imaged with DXA to determine whole-body bone mineral content (BMC), whole-body bone mineral density (BMD), and body composition. The head and the injured femur (or the right femur for uninjured mice) were excluded from the analysis.

### Dynamic Histomorphometry of Bone Formation

Mice were injected with calcein green (10 mg/kg IP; Sigma-Aldrich, St. Louis, MO) and alizarin complexone (30 mg/kg IP; Sigma-Aldrich, St. Louis, MO) 10 days and 3 days prior to death, respectively. After euthanasia, contralateral tibias were collected and embedded in methylmethacrylate plastic (Technovit 7200 and 7210 VLC, EXACT Technologies, Inc., Oklahoma City, OK). Transverse sections were cut on a saw-microtome (Leica 1600SP, North Central Instruments, Inc., Maryland Heights, MO) at the mid-diaphysis to analyze bone formation parameters in cortical bone; frontal sections were cut from the proximal tibia to analyze bone formation parameters in metaphyseal trabecular bone. Slides were initially mounted at a thickness of approximately 600 µm and polished to a thickness of 100 µm. Fluorescent images were obtained on an Olympus IX 70 inverted microscope (Olympus America, Inc., Melville, NY) at 10x magnification, with Nikon NIS Elements imaging software (Nikon Instruments, Inc., Melville, NY), and dynamic histomorphometric analysis was performed using commercial software (Bioquant, Nashville, TN). Mineralizing surface per bone surface (MS/BS), mineral apposition rate (MAR), and bone formation rate per bone surface (BFR/BS) were quantified for the endosteal and periosteal surfaces of cortical bone, and for trabecular bone surfaces.

### Tartrate Resistant Acid Phosphatase Staining of L5 Vertebrae

L5 vertebrae were removed at euthanasia and fixed in 4% paraformaldehyde for 24□hrs. Bones were decalcified in 0.34□M EDTA for 30 days with solution changes every 2–3 days. Vertebrae were embedded in paraffin, frontal sections were taken through the center of the vertebral body at a thickness of 5□µm, and were stained using the TRAP staining system (Sigma-Aldrich, St. Louis, MO) as previously described ^(30)^. Briefly, sections were de-paraffinized, rehydrated, and stained with TRAP working solution with Fast Red Violet as the binding agent at 37°C in the dark until osteoclasts were fully stained. Stained slides were counterstained with 0.02% Fast Green, air dried, and mounted with CoverSafe (American MasterTech, Lodi, CA). Slides were imaged with a Zeiss Axio Imager Z2 Research Microscope (Zeiss United States, San Diego, CA) at 20x magnification. Images were stitched together using Zeiss Zen Pro 2012 Imaging Software. Osteoclast number per bone surface (OcN/BS) and resorbing surface/bone surface (OcS/BS) were calculated using ImageJ ^(31)^.

### ELISA Analysis of Inflammatory Cytokines in Serum

Immediately prior to euthanasia, mice were anesthetized with isoflurane, and a total of 0.5-1 ml of blood was drawn by cardiac puncture and retained for analysis of serum biomarkers. Whole blood serum was centrifuged at 3,000 RPM for 10 minutes, and interleukin-6 (IL-6) and tumor necrosis factor alpha (TNF-α) levels in the supernatant were analyzed using commercially available immunoassay kits, following the manufacturer’s guidelines (R&D Systems Inc., Minneapolis, MN).

## Statistical Analysis

Statistical analyses were performed using JMP Pro 12.0.1 (SAS Institute Inc., Cary, NC). All results are expressed as mean ± standard deviation. Most data were analyzed by three-way analysis of variance (ANOVA) stratified by age (young or middle-aged), days post-injury, and experimental group (Fractured, Sham, or Control) to determine main effects and interactions. DXA data were analyzed using repeated measures ANOVA. Post-hoc analyses were performed using Tukey’s Honest Significant Difference test. Statistically significant differences were identified at p≤0.05. Statistical trends were noted at p≤0.10.

## Results

### Micro-Computed Tomography Analysis of Cortical and Trabecular Bone

Femur fracture resulted in decreased trabecular bone volume and cortical shell thickness of the L5 vertebral body by 2 weeks post-injury in both young and middle-aged mice. By 6 weeks post-injury, microstructural outcomes had fully recovered back to control values in young Fractured mice. However, recovery in middle-aged Fractured mice was unclear; no significant differences were detected between Fractured and Control mice at this time point, but this was primarily due to decreased bone volume in the Control group, rather than recovery in the Fractured group. Sham injury resulted in similar, but lower magnitude, changes in bone microstructure relative to Control mice. For example, at 2 weeks post-fracture we observed 11.2% (p=0.001) and 6.4% (p=0.050) lower BV/TV in young Fractured and Sham mice, respectively, compared to age-matched Control mice, and 18.3% (p=0.035) lower BV/TV in middle-aged Fractured mice compared to Control (Fig. 2). A similar trend was observed at 4 weeks post-injury, although these differences were only statistically significant for middle-aged Fractured vs. Control mice (−19.9%; p=0.010). However, by 6 weeks post-injury, we observed no significant differences between experimental groups for either age group. For young mice, values for Fractured mice increased relative to previous time points to reach Control values. For middle-aged mice, the lack of differences between Fractured and Control mice at this time point was due to an unexpectedly low BV/TV value for Control mice, which was 11-15% lower than week 2 and week 4 Control values, while BV/TV of Fractured middle-aged mice did not change from weeks 2-6. Similar differences were observed for Tb.Th, with 8.5% and 7.2% decreases in young Fractured and middle-aged Fractured mice, respectively, compared to age-matched Controls at 2 weeks post-injury, and subsequent recovery by 4 and 6 weeks post-fracture (Fig. 2).

**Figure 2.**
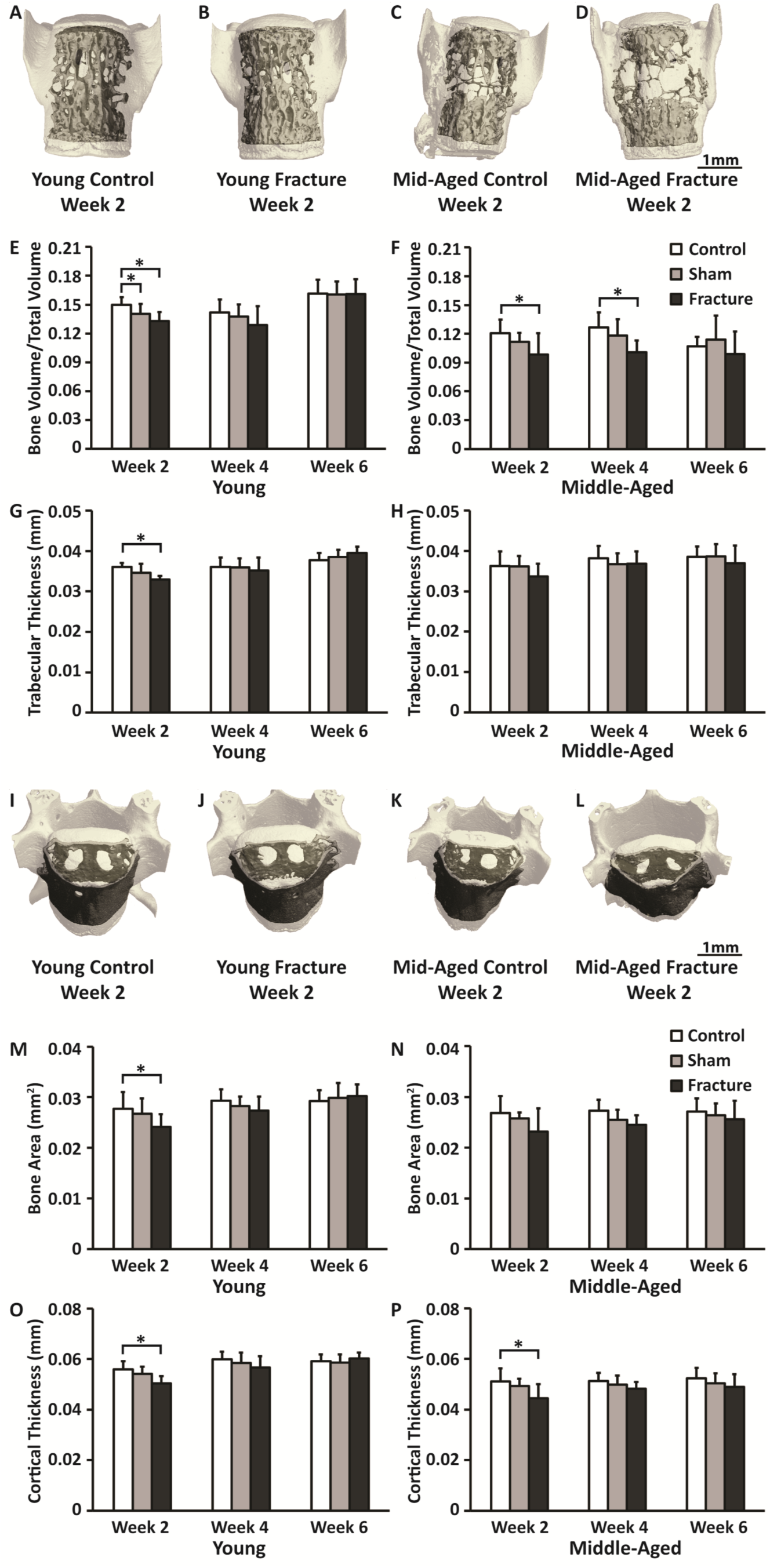
Micro-computed tomography analysis of the trabecular and cortical regions of the L5 vertebral body. (A-D): Representative images of trabecular bone in the L5 vertebral body for (A) young Control, (B) young Fractured, (C) middle-aged Control, and (D) middle-aged Fractured mice at week 2. (E-H): Analysis of trabecular bone volume fraction (E-F) and trabecular thickness (G-H) in young and middle-aged mice. Significant loss of trabecular bone volume and trabecular thickness were observed 2 weeks post-fracture in both age groups, with recovery at later time points. (I-L): Representative images of the cortical shell of the L5 vertebral body for (I) young Control, (J) young Fractured, (K) middle-aged Control, and (L) middle-aged Fractured mice at week 2. (M-P): Analysis of cortical shell bone area (M-N) and cortical thickness (O-P) in young and middle-aged mice. At 2 weeks post-fracture B.Ar. was 12.8% (p=0.027) and 13.6% (p=0.137) lower in young and middle-aged fractured mice, respectively, compared to age-matched control mice. Cortical region analysis of cortical thickness in (O) young and (P) middle-aged mice. Decreased bone area and cortical thickness were observed for both age groups 2 weeks post-fracture, with recovery at later time points. Error bars represent standard deviation. * denotes p≤0.05.

Similar injury-related differences were observed for the cortical shell of the L5 vertebral body (Fig. 2). At 2 weeks post-fracture, B.Ar was 12.8% (p=0.027) and 13.6% (p=0.137) lower in young and middle-aged Fractured mice, respectively, compared to age-matched Control mice. By 6 weeks post-injury, no significant differences were observed between Fractured and Control mice. Similar trends were observed for Ct.Th of the L5 vertebral body cortical shell; 2 weeks post-fracture we observed decreases of 9.7% (p=0.001) and 13.1% (p=0.031) in young and middle-aged Fractured mice, respectively, compared to age-matched Control mice, with no significant differences between injury groups 6 weeks post-injury.

Similar magnitudes of trabecular bone loss were observed in the distal femoral metaphysis of young mice. Young Fractured mice had 11.5% lower BV/TV compared to Control mice 2 weeks post-fracture (p=0.087). However, by 4 and 6 weeks post-fracture no differences were observed between Fractured and Control mice (Supp. Fig. 1). Trabecular bone of middle-aged mice was not measured at the distal femoral metaphysis because there was insufficient trabecular bone for analysis in these mice.

No significant differences were observed in diaphyseal cortical bone parameters as a function of fracture or sham surgery for either young or middle-aged mice (Supp. Fig. 1).

#### Three-Point Bending Mechanical Testing

Three-point bending of both radii and femora revealed significant differences in structural and material properties between young and middle-aged mice, but no differences between injury groups at any time point (Supp. Fig. 2).

### Compression Testing of L5 Vertebral Bodies

No significant differences in structural or material properties of L5 vertebral bodies were observed between the two age groups or between injury groups at any time point (Supp. Fig. 2).

### Finite Element Analysis

Compressive stiffness of L5 vertebral bodies estimated by FE analysis were significantly affected by injury, and followed trends similar to the μCT results for the L5 vertebral body trabecular and cortical shell regions. At 2 weeks post-injury, young and middle-aged Fractured mice exhibited 11.1% (p=0.006) and 19.7% (p=0.016) lower stiffness, respectively, compared to age-matched Control mice (Fig. 3). By 6 weeks post-injury, L5 compressive stiffness of young Fractured mice had fully recovered to Control values.

**Figure 3.**
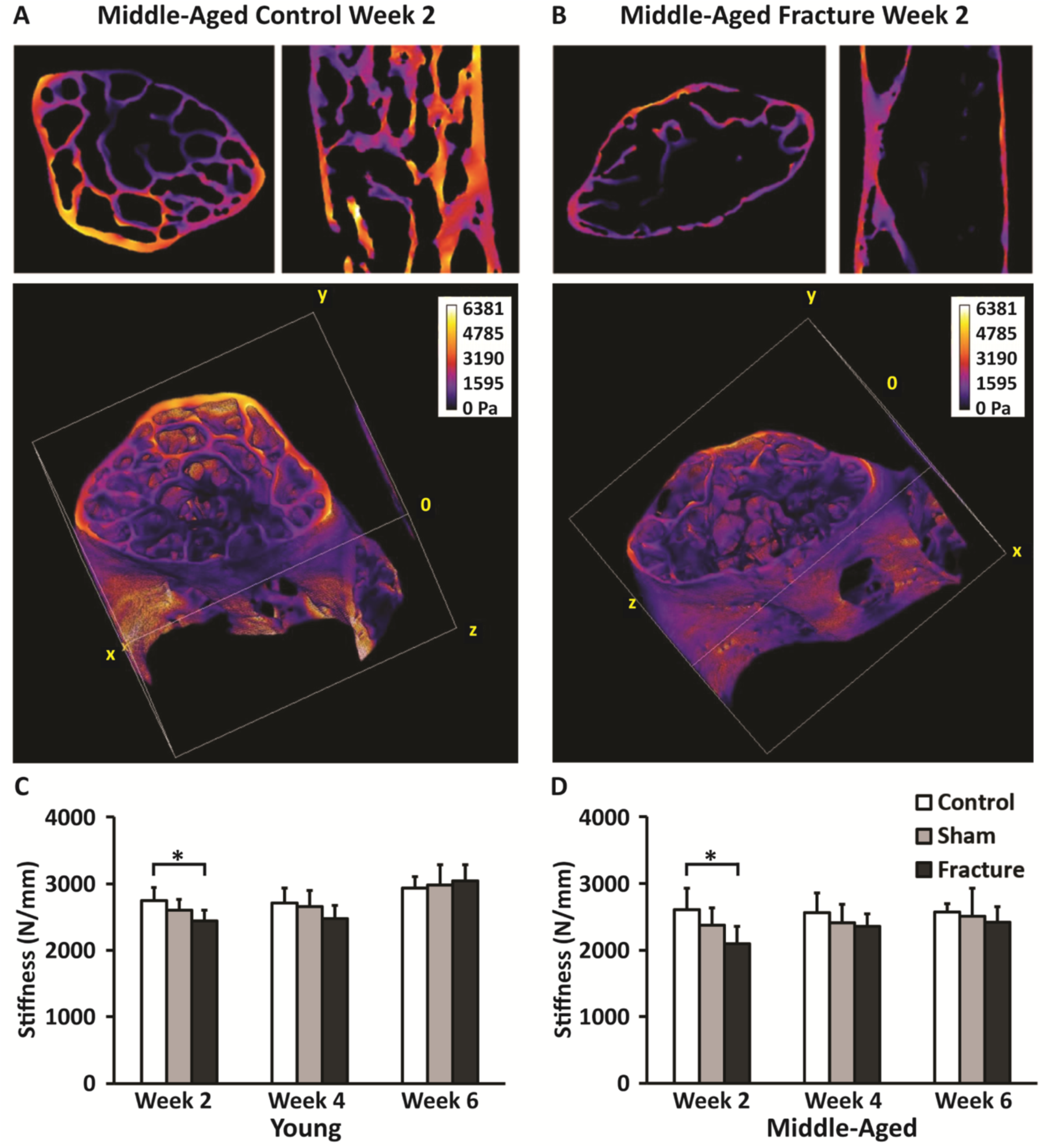
Finite element analysis of the L5 vertebral body. (A-B): Representative images of FE models and Von Mises stress in the longitudinal direction for middle-aged (A) Control and (B) Fractured mice. (C-D): Compressive stiffness estimated by FE analysis in (C) young and (D) middle-aged mice. At 2 weeks post-fracture, both young and middle-aged Fractured mice exhibited significantly lower stiffness than age-matched Control mice. Error bars represent standard deviation. * denotes p≤0.05.

### Quantification of Mouse Voluntary Activity Using Open Field Analysis

Open field analysis revealed significant changes in voluntary movement and activity of Fractured and Sham mice from both age groups 4 days post-injury. At this time point, Total Distance Travelled was decreased in young Fractured mice (−37.0%; p=0.242), middle-aged Fractured mice (−53.6%; p=0.077), and middle-aged Sham mice (−74.8%; p=0.010) relative to age-matched Controls (Fig. 4). At this same time point, Vertical Sensor Breaks were significantly decreased in young Fractured mice (−86.1%; p<0.001), young Sham mice (−50.1%; p=0.011), middle-aged Fractured mice (−81.4%; p<0.001), and middle-aged Sham mice (−69.4%; p<0.001) relative to age-matched Controls (Fig. 4). On day 4 we also observed decreases in Ambulatory Time in young Fractured mice (−41.6%; p=0.071), middle-aged Fractured mice (−49.9%; p=0.005), and middle-aged Sham mice (−57.5%; p<0.001) relative to age-matched Controls (Fig. 4). No significant differences in activity were observed at later time points (weeks 2, 4, and 6 post-injury).

**Figure 4.**
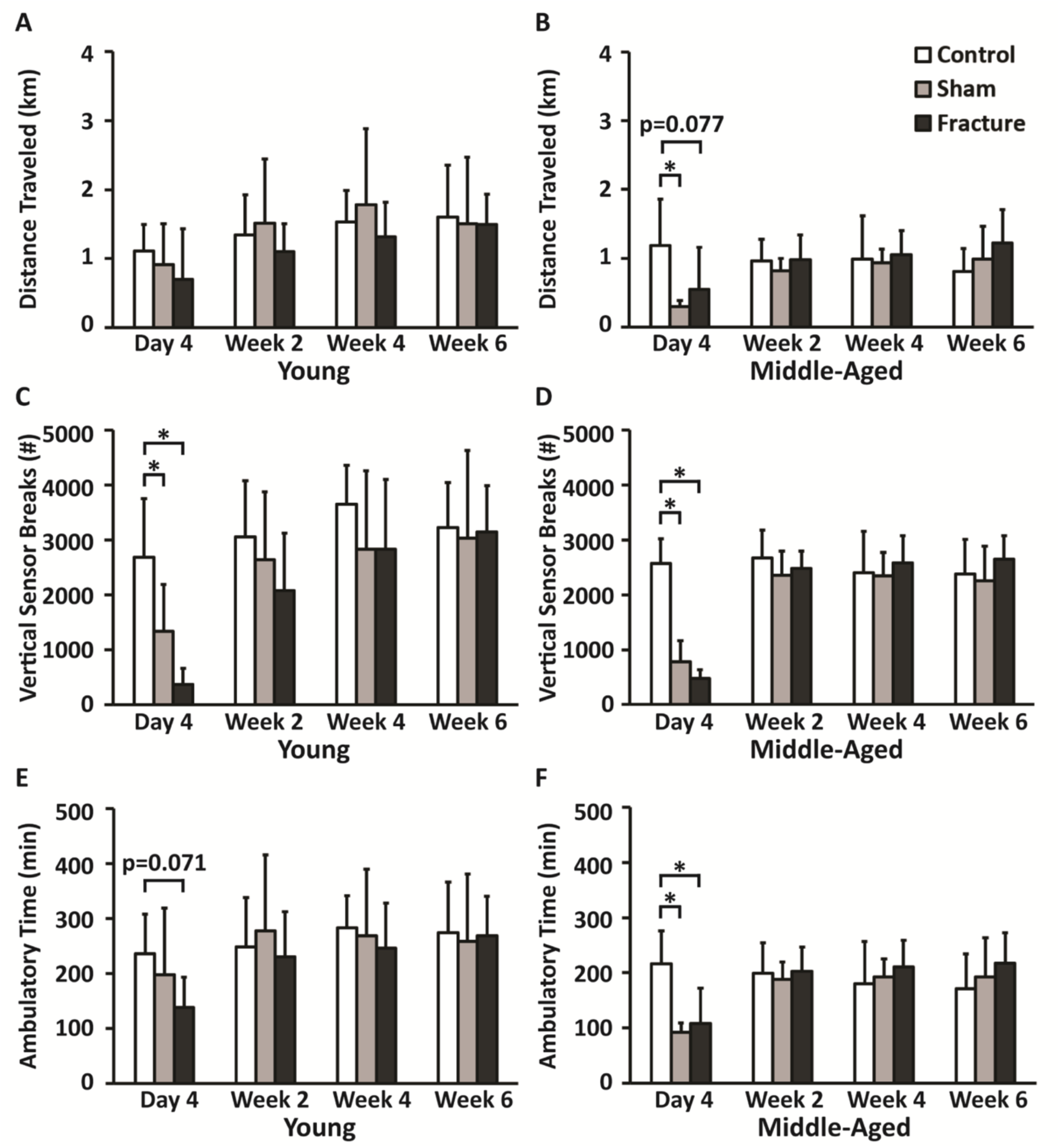
Open field analysis of mouse activity. (A-B): Distance traveled in 24 hours in (A) young and (B) middle-aged mice. On day 4, both young and middle-aged Fractured mice exhibited decreased distance traveled relative to age-matched Controls. (C-D): Vertical sensor breaks in 24 hours in (C) young and (D) middle-aged mice. On day 4, significant decreases in vertical sensor breaks were observed for both Sham and Fractured mice from both age groups relative to age-matched Controls, indicating decreased rearing. (E-F): Ambulatory time in 24 hours in (E) young and (F) middle-aged mice. On day 4, decreases in ambulatory time were observed for Fractured mice from both age groups relative to age-matched Controls. Error bars represent standard deviation. * denotes p≤0.05.

### Dual Energy X-Ray Absorptiometry (DXA)

DXA imaging revealed that both young and middle-aged Fractured mice lost whole-body BMD 2 weeks post-injury (−2.45% for young mice, −1.95% for middle-aged mice) relative to day 4 values (Fig. 5). At subsequent time points, young Fractured mice regained BMD, nearing values for Control mice by 6 weeks post-injury. In contrast, whole-body BMD of middle-aged Fractured mice did not increase between weeks 2-6 post-injury. The time course of BMD changes assessed by DXA was significantly different for young Fractured vs. young Control mice (p=0.029), and for young Fractured vs. middle-aged Fractured mice (p<0.001) (Fig. 5). Similar trends were observed for whole-body BMC, with differences in the time courses between young Fractured vs. young Control mice (p=0.090), and young Fractured vs. middle-aged Fractured mice (p=0.001).

**Figure 5.**
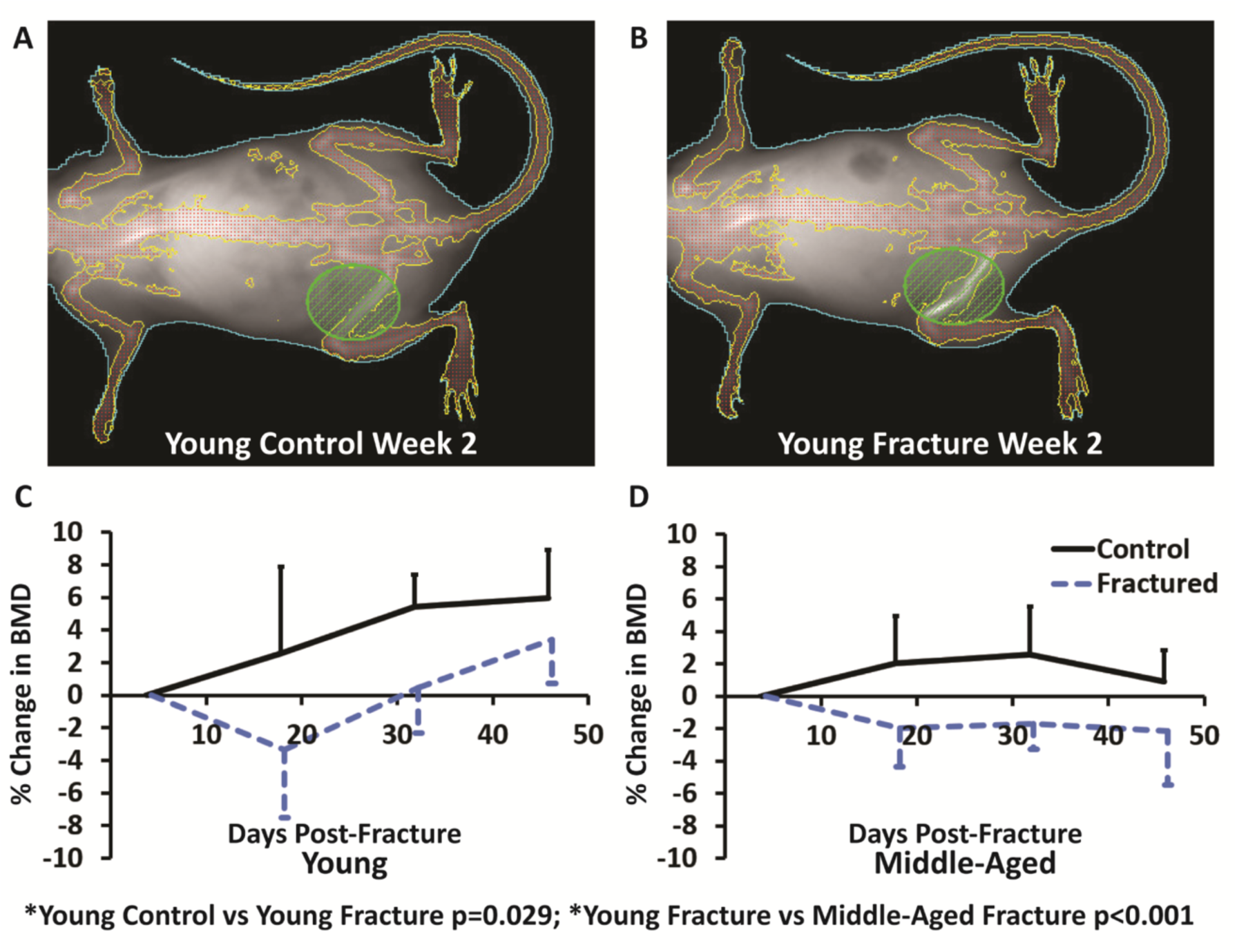
Dual energy x-ray absorptiometry (DXA) imaging of whole-body BMD. (A-B): Representative images of DXA analysis for (A) young Control and (B) young Fractured mice at week 2. (C-D): Percent change in BMD in (C) young and (D) middle-aged Fractured and Control mice. Young and middle-aged Fractured mice lost whole-body BMD 2 weeks post-fracture relative to day 4 values. Young Fractured mice regained BMD at later time points, while Middle-aged Fractured mice did not regain BMD by 6 weeks post-fracture. The time courses of young Fractured vs. middle-aged Fractured mice were significantly different (p<0.001). Error bars represent standard deviation.

### Dynamic Histomorphometry of Bone Formation

Dynamic histomorphometric analysis of bone formation on the endosteal surface of the tibia revealed significant differences between young and middle-age mice for both MS/BS and BFR/BS, but no significant differences between injury groups were observed (Supp. Fig. 3). Little fluorescent labeling was observed on the periosteal surface, with no double labeling on most samples, so MAR and BFR/BS were not calculated for the periosteal surfaces.

Dynamic histomorphometric analysis of trabecular bone of the proximal tibial metaphysis revealed trends toward decreased bone formation in young Fractured mice at 4 and 6 weeks post-injury relative to Control mice (p=0.052). In these mice, BFR/BS was 37.3% lower and 43.7% lower at 4 and 6 weeks, respectively (Fig. 6). No differences were observed at 2 weeks post-injury, and no significant differences in MAR or MS/BS were observed at any time point. Dynamic histomorphometric analysis of trabecular bone was not conducted for middle-aged mice because these mice did not have sufficient metaphyseal trabecular bone for analysis.

**Figure 6.**
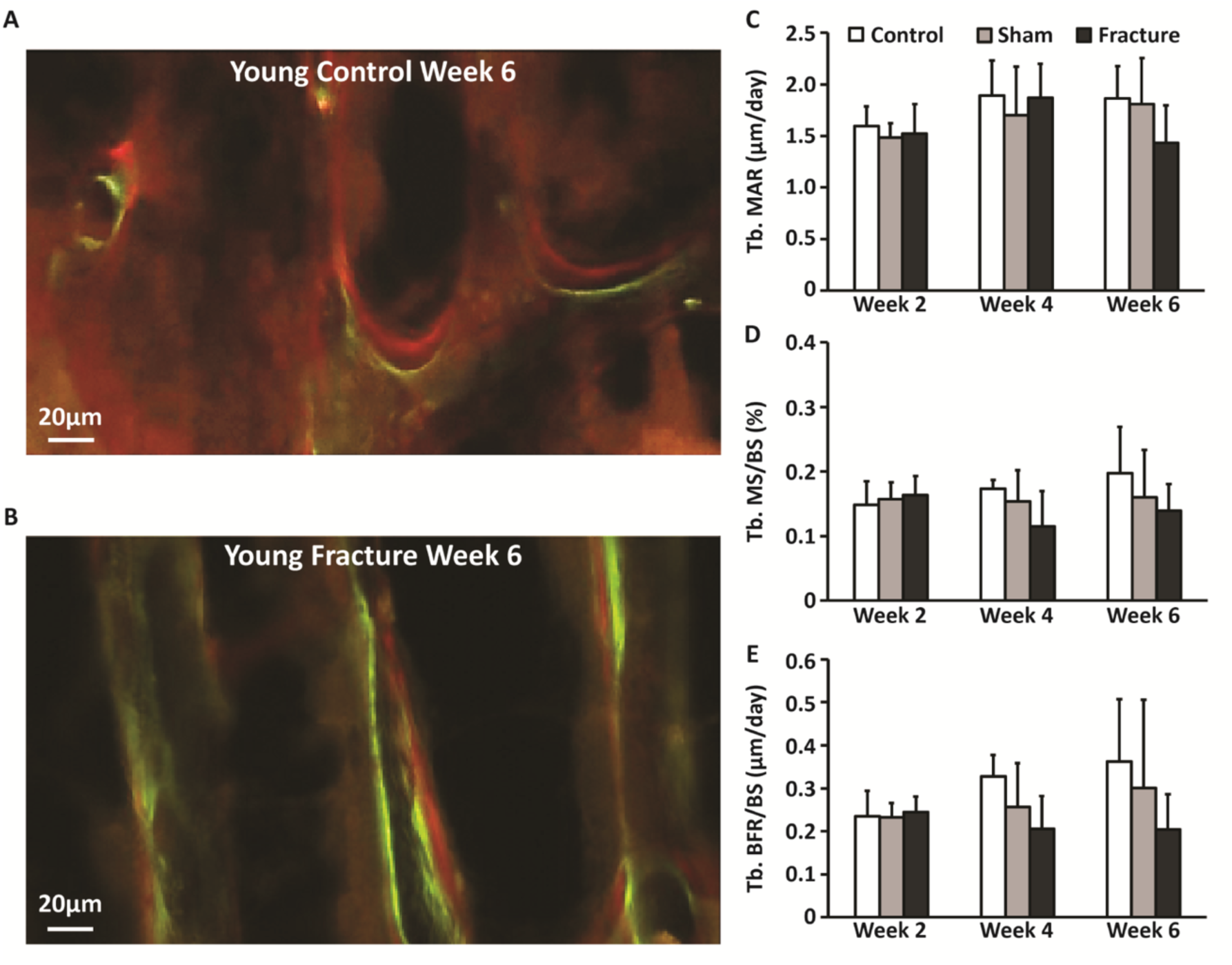
Dynamic histomorphometric analysis of contralateral tibial metaphysis. (A-B): Representative images of frontal sections from (A) young Control and (B) young Fractured mice at week 6. (C-E): Results from young mice for (C) mineral apposition rate, (D) mineralizing surface per bone surface, and (E) bone formation rate per bone surface. Young Fractured mice exhibited trends toward decreased bone formation rate at weeks 4 and 6 post-fracture relative to Control mice (p=0.052). Error bars represent standard deviation.

### Tartrate Resistant Acid Phosphatase Staining

TRAP staining of L5 vertebral bodies revealed significantly increased osteoclast number in trabecular bone of Fractured mice compared to Control mice. On day 3 post-injury, TRAP positive cells per bone surface was increased in both young (+115.9%; p=0.075) and middle-aged (+74.4%; p=0.037) Fractured mice relative to age-matched Control mice (Fig. 7). We also observed a significant main effect of age at day 3, with greater osteoclast number (p=0.003) and resorbing surface (p<0.001) in young mice compared to middle-aged mice (Fig. 7). No significant differences in osteoclast number or resorbing surface were observed at later time points.

**Figure 7.**
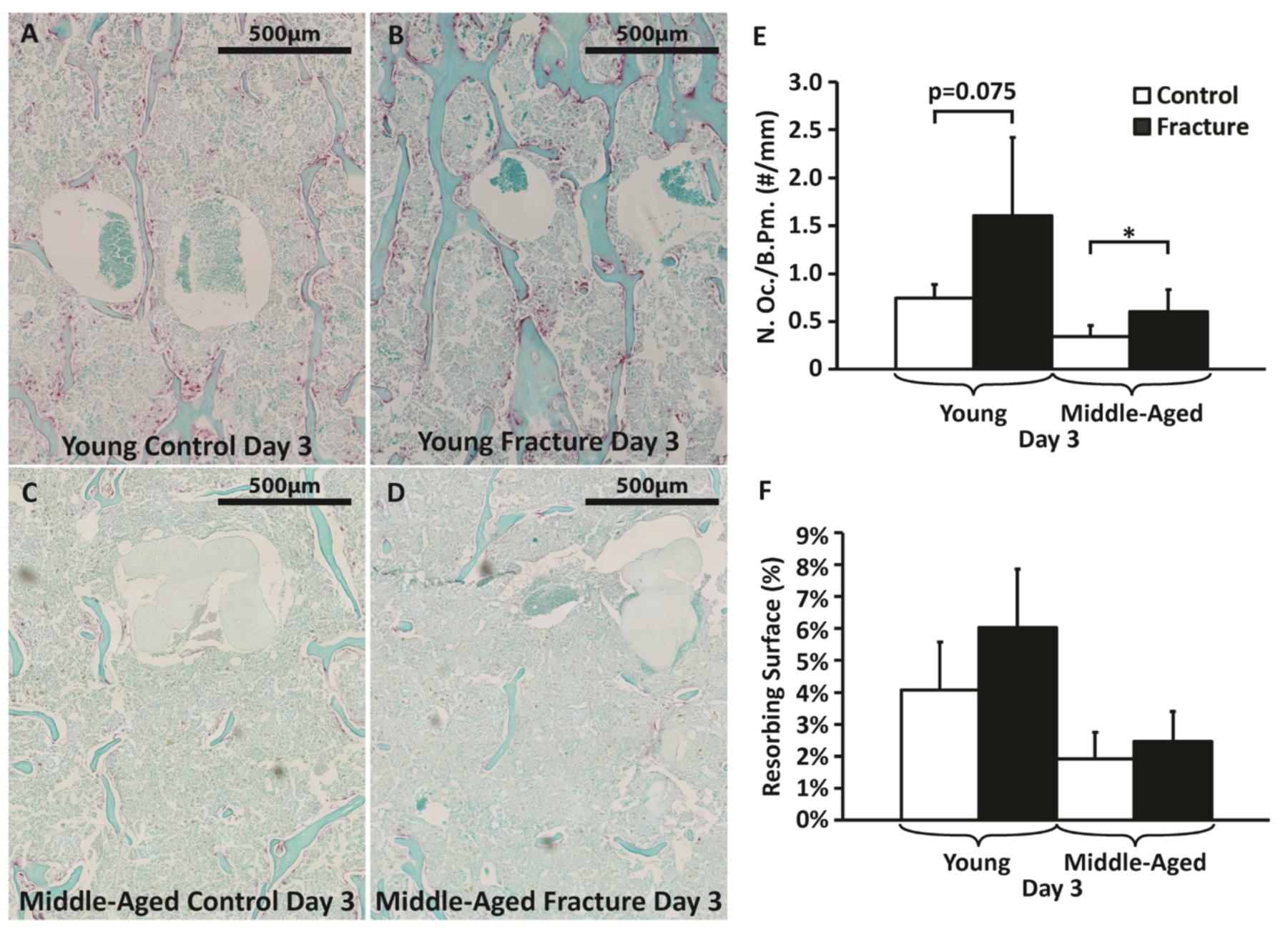
Tartrate-resistant acid phosphatase (TRAP) staining of L5 vertebral body trabecular bone on day 3. (A-D): Representative images of L5 vertebral body trabecular bone from (A) young Control, (B) young Fractured mice, (C) middle-aged Control, and (D) middle-aged Fractured mice. (E-F) Osteoclast number/bone surface and resorbing surface percentage in young and middle-aged mice. On day 3, TRAP positive cells per bone surface were increased in both young and middle-aged Fractured mice relative to age-matched Controls. A main effect of age was observed for both N.Oc./B.Pm. and resorbing surface. Error bars represent standard deviation. * denotes p≤0.05.

### ELISA Analysis of Inflammatory Cytokines in Serum

ELISA analysis of inflammatory cytokines in serum showed significantly increased IL-6 on day 3 post-injury in both young and middle-aged Fractured and Sham mice compared to Control mice. Young Fractured and Sham mice had 3.5-fold (p=0.013) and 6.3-fold (p<0.001) increases, respectively, compared to Control mice, while middle-aged Fractured and Sham mice had 21.9- fold (p<0.001) and 10.2-fold (p<0.001) increases, respectively, compared to Control mice (Fig. 8). No significant differences in IL-6 levels were observed at later time points for either age group. Analysis of serum TNF-α showed no differences in cytokine levels in either age group at any of the time points, and in fact was not detectable above background values for most serum samples.

**Figure 8.**
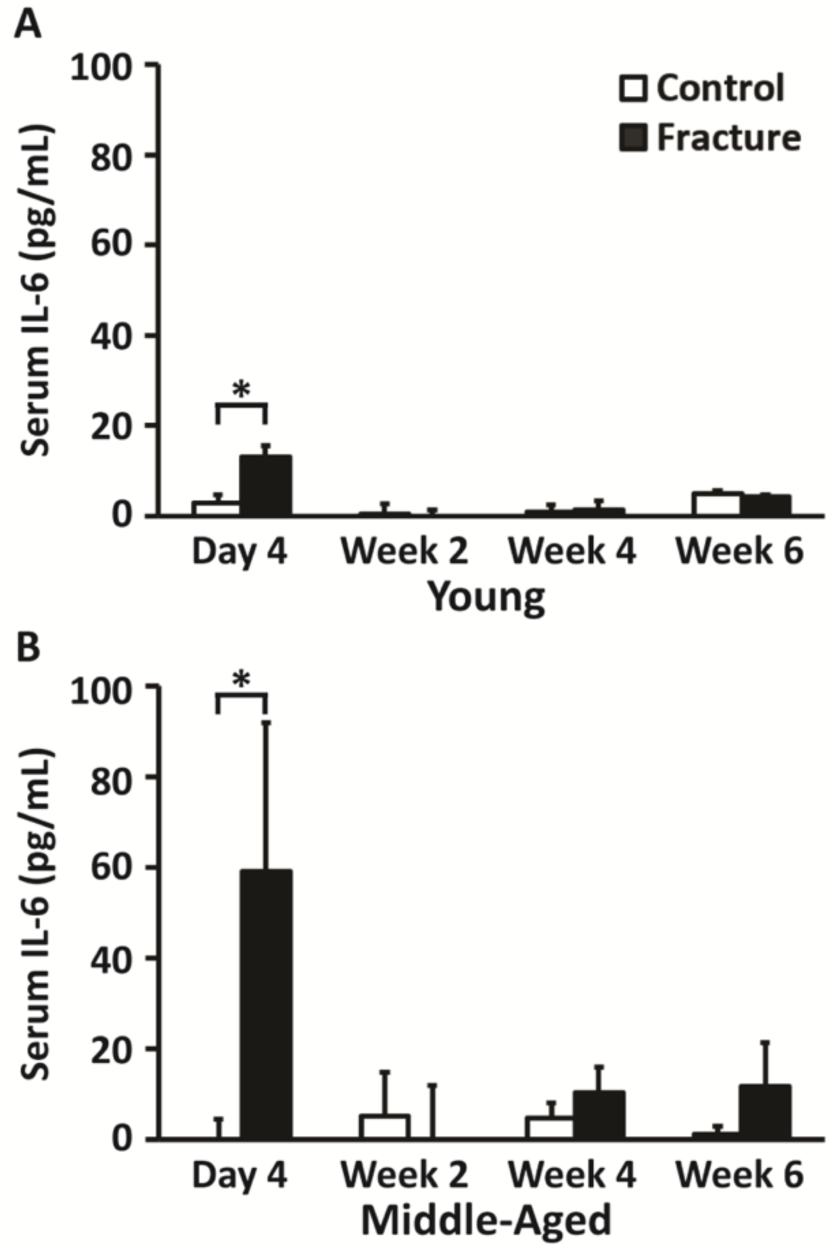
Enzyme-linked immunosorbent assay (ELISA) of serum interleukin-6 levels in (A) young and (B) middle-aged mice. At day 3, young Fractured mice had a 3.5 fold increase in serum IL-6, while middle-aged Fractured mice had a 21.9 fold (p<0.001) increase in serum IL-6 compared to age-matched Control mice. Error bars represent standard deviation. * denotes p≤0.05.

## Discussion

In this study, bone loss and recovery following fracture in young and middle-aged mice were measured. Consistent with our initial hypothesis, we found that femur fracture resulted in loss of trabecular and cortical bone volume in the L5 vertebral body and decreases in whole-body BMD assessed by DXA 2 weeks post-injury. This early bone loss was accompanied by increased levels of inflammatory cytokine in serum, increased osteoclast number and activity, and changes in voluntary movement of mice at early time points post-fracture. At later time points we observed an age-dependent recovery from this initial bone loss, with young mice exhibiting a full recovery of bone microstructure by 6 weeks post-fracture, while middle-aged mice either did not show any recovery (DXA) or exhibited partial recovery (µCT) of bone 6 weeks post-fracture. These data establish a systemic bone loss response following fracture that actively reduces the bone mass and strength of the skeleton; this phenomenon may partially explain why an initial fracture increases the risk of future fractures in human subjects.

The trabecular bone loss at the L5 vertebral body 2 weeks after femoral fracture observed in this study is consistent with our previous studies that showed loss of trabecular bone from the contralateral femoral epiphysis and L5 vertebral body following non-invasive injury of the anterior cruciate ligament in mice ^(32,33)^. It is also partially consistent with the previously described regional acceleratory phenomenon (RAP) ^(13–15)^ and systemic acceleratory phenomenon (SAP) ^(16,17)^ that have been reported in bone after a fracture or other insult to the skeleton. These injuries initiated increased bone formation and resorption rates near the site of injury (RAP) and at distant skeletal sites (SAP). In our study, we found that femur fracture increased osteoclast number and activity in the trabecular bone of the L5 vertebral body 3 days post-fracture. However, we also found that bone formation in the trabecular compartment of the proximal tibial metaphysis was not changed 2 weeks post-fracture, and trended toward decreased bone formation rates at 4 and 6 weeks post-fracture, rather than exhibiting increased rates following fracture as expected. It is unclear why these disparate outcomes were observed, but it may relate to factors such as the animal model, injury type, skeletal site, or time points analyzed.

Age is an important factor affecting bone metabolism and bone adaptation to injury or mechanical loading. Older animals generally exhibit a diminished anabolic response to mechanical loading ^(34–38)^ and a diminished catabolic response to mechanical unloading ^(39)^ relative to younger animals. This reduced mechanosensitivity in older animals suggests that post-fracture bone loss in middle-aged mice may be partially due to non-mechanical factors such as systemic inflammation. This may also support the possible age-related differences in recovery from systemic bone loss. Aging has also been shown to hinder fracture healing and increase the inflammatory response to injury ^(40–43)^, with aged mice exhibiting increased fracture callus size and a delayed healing response to fracture ^(40)^. Our quantification of serum inflammatory cytokines is in agreement with these previous studies. We observed significant increases in serum concentrations of IL-6 at 3 days post-fracture in both young and middle-aged mice, but this increase was considerably greater in middle-aged mice. Age-related differences in systemic bone loss and recovery are also supported by clinical data showing that fractures between the ages of 20-50 are predictive of future fracture risk in women, while fractures before age 20 are not correlated with future fractures ^(3)^.

Underlying mechanisms driving the systemic bone loss response have not been established, but likely include injury-induced systemic inflammation and altered mechanical loading (cage activity) after injury. Following fracture we observed a considerable increase in serum levels of IL-6, particularly in middle-aged mice. Inflammatory cytokines such as interleukin 1 (IL-1), interleukin 6 (IL-6), and tumor necrosis factor (TNF) are elevated in inflammatory states, and have the ability to modulate bone mass by directly or indirectly promoting osteoclastogenesis ^(44,45)^. IL-1 plays an important role in early response to fracture ^(46,47)^, and also stimulates expression of IL-6 by osteoblasts ^(48,49)^. IL-6 stimulates production of RANK ligand by osteoblasts, which catalyzes the formation of osteoclasts ^(50)^; IL-6 also induces the differentiation of pre-osteoblasts ^(51–53)^. The effect of acute injury-induced inflammation on the systemic proliferation of osteoclasts is not as well established, but it is possible that increased levels of inflammatory cytokines directly contributed to the increased osteoclast number and increased bone resorption we observed in the lumbar spine 3 days post-fracture. We also observed some significant changes in the voluntary cage activity (mechanical loading) of mice 4 days post-fracture. At this time point, Fractured mice tended to walk less overall, with considerably decreased incidence of rearing. These changes in cage activity following fracture may have contributed to systemic bone loss, as the association between disuse and bone resorption is well established. However, disuse following injury is a very clinically relevant occurrence that would likely affect human patients as much or even more than mice. In this way, we consider decreased voluntary movement to be one of several relevant factors contributing to systemic bone loss.

From an evolutionary perspective, systemic bone loss following fracture may be a mechanism of mineral utilization for fracture callus formation. One of the primary functions of bone is as a reservoir for mineral; utilization of this mineral to address an acute need (fracture callus formation) may be sufficient to cause lasting effects on fracture risk at other skeletal sites if bone mass is not fully recovered. This is supported by a previous animal study that showed that calcium supplementation improved fracture healing in osteoporotic rats ^(54)^. Similarly, a recent study by Fischer et al. ^(55)^ reported that fracture healing was marginally disturbed in Calcium/Vitamin D-deficient mice, and that deficient mice displayed significant systemic osteoclast activity and reduced bone mass in the intact skeleton post-fracture, suggesting considerable calcium mobilization systemically during bone healing. This study also found that Ca/VitD supplementation post-fracture improved bone healing and prevented systemic bone loss by reducing bone resorption.

Many epidemiological studies have shown increased fracture risk after an initial fracture ^(1–10)^, but it has not yet been established that systemic bone loss following fracture results in a deficit in bone strength or other mechanical properties. In the current study, mechanical testing of mouse bones did not show any significant differences in mechanical properties as a function of injury. This could be largely due to the fact that it is technically challenging to perform mechanical testing of small bones with high fidelity, and errors associated with processing, alignment, or loading of bone specimen may have introduced considerable variability in the data ^(56)^. However, FE analysis of L5 vertebral bodies, a more controlled method for assessing mechanical outcomes, predicted significant differences in compressive stiffness in Control versus Fractured mice. Differences between injury groups using this method were observed at early time points, and followed similar trends to the µCT from the lumbar spine. Additionally, the magnitude of bone loss observed using µCT data was relatively small (typically 10-20% decrease in trabecular bone volume fraction), so it is possible that these differences also translate to small differences in mechanical properties that can be difficult to detect. However, it has been shown that small differences in trabecular bone volume can translate to much larger changes in mechanical strength ^(57)^, therefore it is possible that even small amounts of systemic bone loss could translate to significant increases in fracture risk, especially when accumulated across a large population.

This study had several notable strengths. We clearly established that the systemic bone loss phenomenon occurs in mice using both whole-body analysis (DXA) and microstructural analysis of skeletal sites distant from the injury (µCT). We also quantified biological and mechanical factors that could contribute to systemic bone loss following fracture. We investigated the effect of age on systemic loss and recovery of bone, and age-related differences were observed in both biological and mechanical outcomes. Future studies could further investigate systemic bone loss in even older animals or ovariectomized mice (representative of post-menopausal women). The study also had some important limitations. Femur fracture resulted in changes in overall cage activity, but also likely altered gait patterns that may cause compensatory loading of other limbs. We evaluated an axial skeletal site (L5 vertebral body) to try to minimize the effect of altered gait patterns, but the overall effect of asymmetric gait is not known. The mechanical testing data in this study is also a limitation because of the technical difficulties and high level of variance associated with processing and testing of small specimens. These limitations may have obscured mechanical outcomes assessed by mechanical testing, especially if the effect size was small. Finite element analysis provided complimentary data that more precisely determined mechanical properties of the L5 vertebral body, but this analysis used linear elastic inputs, which is a simplification of the actual mechanical behavior of bone. Another limitation is that many of our outcomes were cross-sectional. For these data, changes over time were iterated from different groups of mice, which potentially adds variability when making comparison across experimental groups. We also used different skeletal sites for different analyses (i.e., L5 vertebral body for analysis of bone resorption, proximal tibia for analysis of bone formation). This requires an assumption that bone turnover is similar at these different skeletal sites, but it is currently unknown if this assumption is valid. Another limitation is that middle-aged female mice did not have sufficient trabecular bone in the tibial or femoral metaphysis, therefore analysis of microstructure and bone formation at these sites was not possible. Finally, we detected differences in both systemic inflammation and voluntary cage activity at early time points following fracture. It is likely that both of these contribute to systemic bone loss, but the individual contributions of each of these mechanisms have not been established. It is also possible that additional mechanisms that have not yet been identified also contribute to systemic bone loss. Further investigation of the injury-induced response is needed to further characterize mechanisms contributing to bone remodeling following fracture.

## Conclusions

In this study, we showed that femur fracture initiated a systemic bone loss response in mice, and that there were significant age-related differences in this response. Bone fracture initiated injury-induced systemic inflammation and decreased mechanical loading, both of which can modulate the function of bone cells and negatively affect both bone quantity and bone quality. Importantly, these injury-induced responses are also operative in human subjects following traumatic fracture, so it is reasonable to predict that this systemic bone loss response may meaningfully contribute to fracture risk in osteoporotic subjects. Uncovering the etiology of this phenomenon will allow us to inform treatments aimed at preserving lifelong skeletal health for the aging population.

## Acknowledgments

We would like to thank Todd Tolentino, Heather Tolentino, and Samrrah Raouf at the UC Davis Mouse Biology Program – MMPC Energy Balance, Exercise & Behavior Core (NIH grant U24-DK092993) for performing the Open Field and DXA analyses. We would also like to thank Cristal Yee, Alfred Li, and Tamara Alliston at the University of California San Francisco – Skeletal Biology Core for performing TRAP staining and imaging.

Research reported in this publication was supported by the American Society for Bone and Mineral Research Junior Faculty Osteoporosis Research Award, by the UC Davis Clinical and Translational Science Center, and by the National Institute of Arthritis and Musculoskeletal and Skin Diseases, part of the National Institutes of Health, under Award Number AR071459. The content is solely the responsibility of the authors. The funding bodies were not involved with design, collection, analysis, or interpretation of data; or in the writing of the manuscript. The authors have no conflicts of interest to disclose.

---

Supplemental Figure 1. Micro-computed tomography results of the trabecular and cortical regions of the contralateral femur and the contralateral radius cortical region. (A-B): Trabecular bone volume fraction and trabecular thickness of the distal femoral metaphysis trabecular region. A trend toward decreased trabecular bone volume was observed in young Fractured mice compared to Control mice at week 2. (C-F): Bone area and cortical thickness of the mid-diaphysis of contralateral femurs from young and middle-aged mice. (G-J): Bone area and cortical thickness of the mid-diaphysis of contralateral radii from young and middle-aged mice. No differences in cortical bone microstructure were observed as a consequence of fracture for either age group. Error bars represent standard deviation.

Supplemental Figure 2. Mechanical testing of femurs, radii, and vertebral bodies. (A-B): Stiffness of contralateral femurs tested in three-point bending. (C-D): Stiffness of contralateral radii tested in three-point bending. (E-F) Stiffness of L5 vertebral bodies tested in compression. Error bars represent standard deviation.

Supplemental Figure 3. Dynamic histomorphometric analysis of bone formation on the cortical surface of the contralateral tibial diaphysis. (A-D): Representative images of contralateral tibia cross-sectional images of (A) young Control, (B) young Fractured, (C) middle-aged Control, and (D) middle-aged Fractured mice. (E-F): Bone formation rates on the endosteal surface of (E) young and (F) middle-aged mice. No differences were observed between injury groups for MAR, MS/BS, or BFR/BS on the endosteal surface. MAR and BFR/BS were not calculated on the periosteal surface because of insufficient double labeling. Error bars represent standard deviation.

